# Identification of Functional NOD2-Specific Aptamers Using APT-SNAP Microarrays to Screen Zebrafish Nod-like Receptors

**DOI:** 10.64898/2026.11.24.768724

**Authors:** Paige E. Messa, Keenan S. Pearson, Brian J. Conti, Daniel T. Vogt, Noah R. Nicol, Vaishnav Kumar, Elise S. Ozers, Christopher L. Warren, Mary S. Ozers

## Abstract

NOD-like receptors (NLRs) are intracellular immune sensors that monitor for pathogen- and damage-associated molecular patterns (PAMPs, DAMPs) and initiate innate immune signaling pathways upon detection. Dysregulation of NLR signaling contributes to numerous inflammatory diseases and cancers, yet mechanistic studies of NLR proteins are limited by the availability of reliable detection reagents, particularly in emerging model organisms such as zebrafish (*Danio rerio*). Although zebrafish provide a powerful system for studying innate immunity, few validated antibodies exist for zebrafish NLR proteins. Here, we describe the development of aptamer-based detection reagents targeting zebrafish NLRs using the APT-SNAP platform, a high-density aptamer microarray technology that enables rapid identification of protein-binding nucleic acid ligands. Using this approach, we identified multiple DNA aptamers that bind zebrafish NLR proteins and selected candidates targeting NOD2 for further characterization. Orthogonal validation assays, which include enzyme-linked apta-sorbent assays (ELASAs), pull-downs, biolayer interferometry (BLI), and fluorescence microscopy, confirmed that the lead aptamers bind zebrafish NOD2 with high affinity and specificity, and function effectively in biochemical and cellular assays. These results demonstrate that APT-SNAP generated validated, native-state affinity reagents for a zebrafish immune protein where commercial antibodies failed, providing new molecular tools for investigating NLR biology in zebrafish. More broadly, this strategy establishes a generalizable approach for generating affinity reagents against challenging protein targets that lack reliable antibodies.

## INTRODUCTION

NOD-like receptors (NLRs) are a key component of the innate immune system that sense and coordinate cellular responses to pathogens and other forms of cellular stress. Disruptions in NLR function are implicated in a variety of inflammatory and autoimmune disorders, including inflammatory bowel disease (IBD) (1–4), type 1 and 2 diabetes (T1D, T2D) (5, 6), asthma (7), and several cancers (8). Consequently, NLR signaling pathways are an important focus of both basic research and therapeutic development. Despite their biological significance, the study of NLR proteins remains challenging due to their complex functional mechanisms, differences between model systems, and the limited availability of reliable detection reagents.

NLRs function as cytosolic pattern recognition receptors that detect pathogen-associated molecular patterns (PAMPs) and damage-associated molecular patterns (DAMPs), which are endogenous signals indicative of cellular stress or injury (8, 9). Structurally, NLRs typically contain a central nucleotide-binding oligomerization domain (NACHT), a C-terminal leucine-rich repeat domain involved in ligand recognition, and a variable N-terminal effector domain that mediates interactions with adaptor proteins (Fig. 1A) (9). Based on their N-terminal domains, NLRs are further divided into subfamilies that dictate their downstream signaling functions (10, 11). Upon activation, NLRs oligomerize to form multiprotein complexes or inflammasomes that initiate signaling events including caspase activation and cytokine release, pyroptosis, and activation of NF-κB and MAPK pathways, ultimately coordinating inflammatory and immune responses (8). The diversity of stimuli and the overlapping functions of NLR family members complicate mechanistic studies of these receptors.

**Figure 1.**
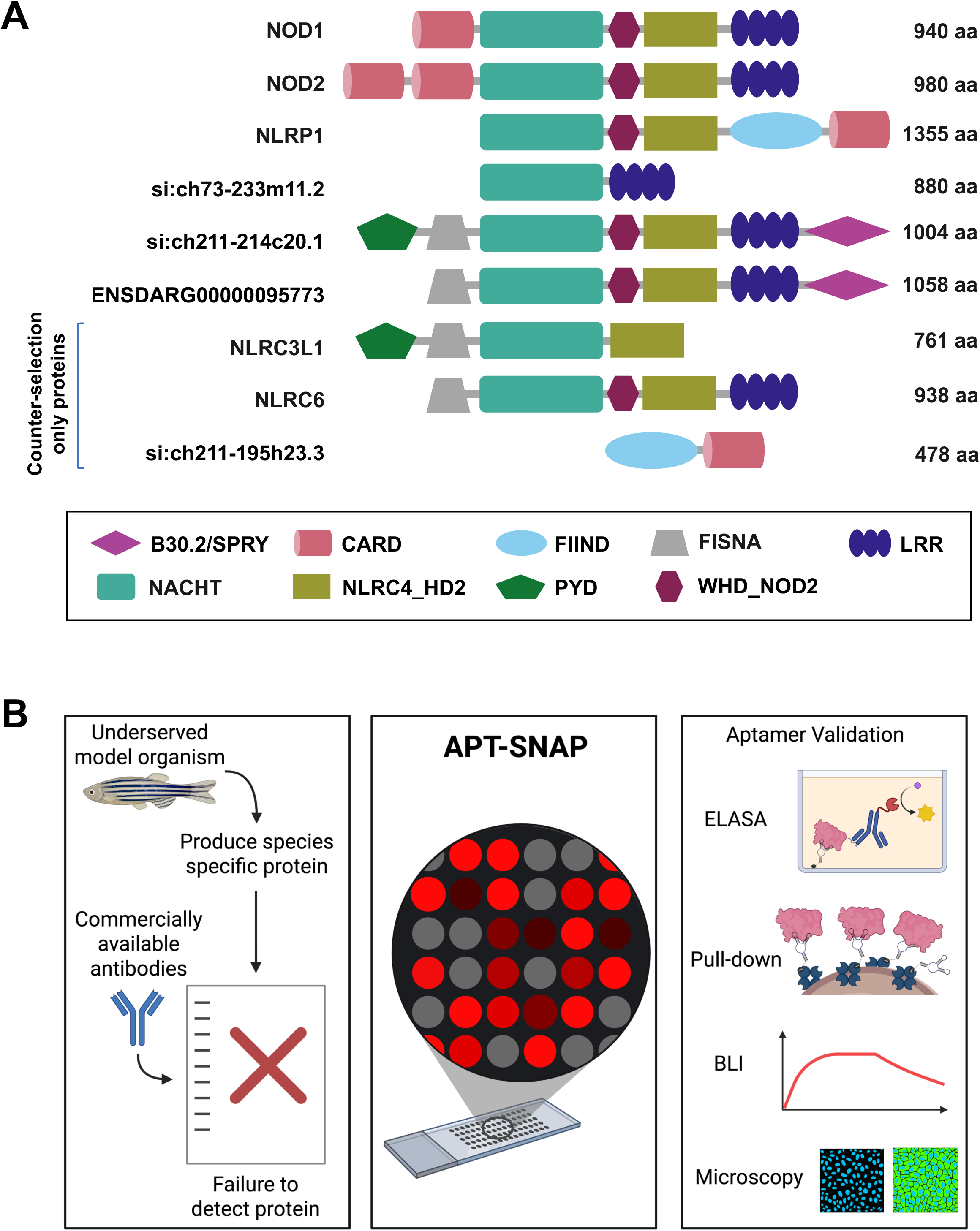
Experimental workflow and domain architecture of zebrafish NLR proteins analyzed in this study. A) Domain organization of zebrafish NLR target and counter-screening proteins used in this study with total amino acid (aa) lengths indicated. Domains include B30.2/SPRY (protein-protein interaction domain), CARD (caspase recruitment domain), FIIND (function-to-find domain), FISNA (fish-specific NACHT associated domain), LRR (leucine-rich repeat domain), NACHT (nucleotide-binding domain), NLRC4_HD2 (NLRC4 helical domain HD2), PYD (pyrin domain), and WHD_NOD2 (NOD2 winged helix domain). Counter-selection proteins used only during APT-SNAP screening are indicated. B) Overview of the APT-SNAP aptamer discovery and validation workflow. Recombinant zebrafish NLR proteins were expressed and purified, screened against a high-density aptamer array using the APT-SNAP platform, and candidate aptamers were subsequently validated using ELASA, pull-down, biolayer interferometry (BLI), and fluorescence microscopy assays.

The zebrafish (*Danio rerio*) has emerged as a powerful model organism for studying innate immunity. Zebrafish are well suited for high-throughput experimentation due to their low cost, genetic tractability, and optical accessibility (12). Of note, zebrafish larvae lack a fully mature adaptive immune system until approximately four weeks after fertilization, allowing innate immune responses to be studied in relative isolation (13). Comparative genomic analyses have revealed substantial conservation between human and zebrafish immune pathways (14, 15), including highly conserved NLR family members such as NOD1 and NOD2 that are two central regulators of bacterial peptidoglycan sensing. At the same time, zebrafish possess a large expansion of NLR genes resulting from duplication events, including approximately 368 additional NLR-like proteins with limited homology to mammalian counterparts (16). While zebrafish studies have provided significant insight into other classes of pattern recognition receptors such as Toll-like receptors (TLRs) (17), comparatively little work has been done to investigate NLR biology in this model system.

A major barrier to studying zebrafish NLRs is the lack of reliable protein detection reagents. Although antibodies exist for many mammalian NLR proteins, few have been validated for recognition of zebrafish orthologs, and commercial antibodies specific to zebrafish NLRs are largely unavailable (18, 19). This scarcity of reagents has limited the ability to investigate NLR expression, localization, and function in zebrafish.

Aptamers are an attractive alternative to antibodies for protein detection and targeting applications. Aptamers are short nucleic acid-based reagents capable of binding protein targets with high affinity and specificity (20, 21). These molecules can recognize both cell-surface and intracellular targets and can be selected for specific localization or biological activity (22–26). In many cases, aptamers exhibit sub-nanomolar dissociation constants and can discriminate between closely related proteins (22, 27). Because aptamers are nucleic acid-based molecules, they can be synthesized inexpensively, exhibit consistency between batches/manufacturers, are thermally stable, and can be developed entirely in vitro without the use of animals.

Here, we describe the development and validation of aptameric detection reagents targeting zebrafish NLR proteins using the APT-SNAP platform (Fig. 1B), a novel high-density aptamer microarray technology we developed that enables rapid identification of high-affinity protein-binding aptamers without iterative enrichment steps. Using this approach, we identified a suite of aptamer candidates against zebrafish NLR targets. We then performed orthogonal validation of the highest-performing candidates against zebrafish NOD2. These results provide new molecular tools for studying NLR biology in zebrafish and establish a generalizable strategy for developing affinity reagents for challenging targets that lack reliable antibodies.

## RESULTS

### Expression and purification of zebrafish NLR proteins using a baculovirus expression system

Six members of the zebrafish NLR family were selected as targets for aptamer discovery, including three proteins with strong structural and functional conservation to human orthologs (NOD1, NOD2, and NLRP1), one protein with partial similarity to human NLRP14 (si:ch73-233m11.2), and two proteins from the fish-specific NLR multigene family (si:ch211-214c20.1 and ENSDARG00000095773). Three additional proteins (NLRC3L1, NLRC6, and si:ch211-195h23.3) were included as counter-selection proteins to ensure that each Pfam domain represented among the targets was present in at least two proteins (Fig. 1A and Table S1).

Recombinant zebrafish NLR proteins containing N-terminal His-tags were expressed in insect cells using a baculovirus expression system and purified by immobilized metal affinity chromatography. Protein purity and potential degradation were evaluated by SDS-PAGE followed by Coomassie staining (Fig. 2A). Expression was further confirmed by western blot using an anti-His-tag antibody, which detected bands corresponding to the expected molecular weights of the recombinant proteins (Fig. 2B and Table S1). Mass spectrometry analysis confirmed the identity of the proteins present in these bands (data not shown).

**Figure 2.**
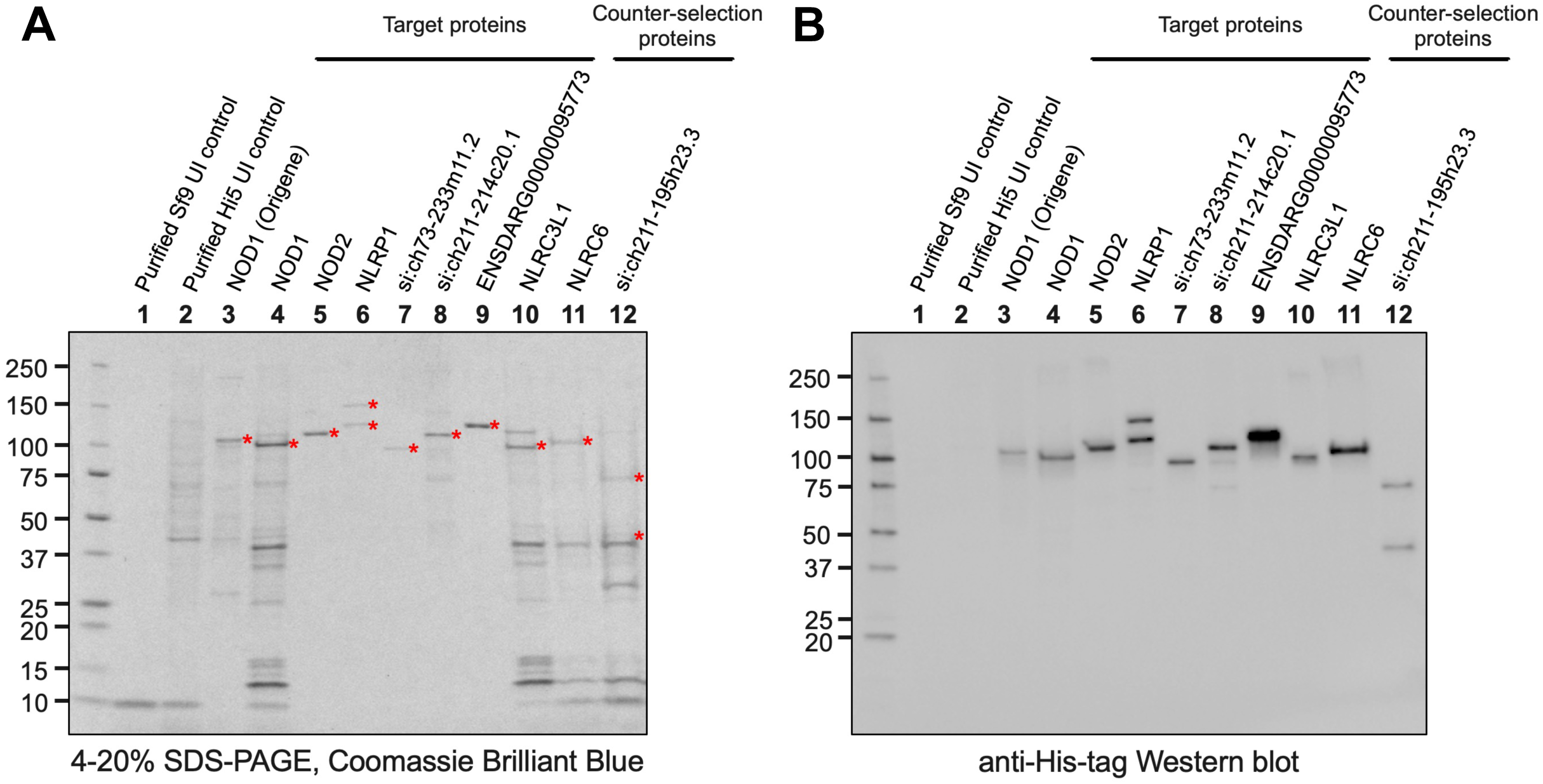
Expression and purification of zebrafish NLR proteins. A) SDS-PAGE analysis of purified NLR proteins expressed in ExpiSf9 or Hi5 insect cells using a baculovirus expression vector system. Purified Sf9 control and purified Hi5 control represent uninfected (UI) cell controls. Molecular weight markers are shown on the left. Target proteins and counter-selection proteins are indicated above the gel. B) Western blot analysis of purified NLR proteins using an anti-His tag antibody. Samples correspond to those shown in panel A. Molecular weight markers are indicated on the left, with additional detail provided (Table S1).

For NLRP1 and the counter-selection protein si:ch211-195h23.3, western blot analysis revealed an additional lower-molecular weight band. Mass spectrometry analysis identified these bands as truncated versions of the proteins (data not shown). This is likely an expected autoproteolytic event associated with the function-to-find (FIIND) domain present in these two NLR proteins (Fig. 1A) (28), indicating a known cleavage of these proteins rather than degradation.

NOD1 was expressed but exhibited low solubility and limited yield in the baculovirus system (Fig. 2A). During this study, the original zebrafish NOD1 entry in the UniProt database was updated following annotation of an alternative transcript with domain architecture more consistent with canonical NLR proteins. Because of the limited yield of the original construct, a recombinant NOD1 protein corresponding to the updated sequence was subsequently obtained from a commercial source (Origene) for downstream analyses.

### Commercial antibodies against mammalian NLRs do not recognize zebrafish orthologs

After establishing recombinant expression of zebrafish NLR proteins, we next evaluated whether commercially available antibodies raised against mammalian NLR proteins could be used to detect these orthologs. Although many NLR proteins are conserved between human and zebrafish, sequence identity between orthologs can vary substantially, which may limit the utility of antibodies developed against mammalian proteins for detecting zebrafish counterparts. The percent identities for zebrafish and human NOD1, NOD2, and NLRP1 are 51.85%, 47.08%, and 28.63%, respectively, despite their conserved functionality, as determined by NCBI BLASTp. To determine whether existing antibodies could be used to detect zebrafish NLR proteins examined in this study, commercially available antibodies raised against human NOD1, NOD2, NLRP1 and NLRP14 (homologous to si:ch73-233m11.2) were evaluated by western blot analysis.

Antibodies were tested using lysates from recombinant baculovirus infected insect cells overexpressing the corresponding zebrafish NLR proteins alongside mammalian cell lysates expressing the human orthologs as positive controls (obtained from Origene). Detection of FLAG- or His-tagged proteins served as controls for successful protein expression. Antibody performance was evaluated based on the presence of a band at the expected molecular weight for each target protein. The results of these experiments are summarized in Tables 1, S2, S3, and S4.

**Table 1.** Human NOD2 antibody testing summary.

| **Antibody** | **Species Reactivity** | **Immunogen** | **Detects Human NOD2?** | **Detects Zebrafish NOD2?** |
|---|---|---|---|---|
| PA5-104317 | Human, Mouse | Synthetic peptide derived from human NOD2 corresponding to a region within N-terminal amino acids | Yes | No |
| 14-5858-81 | Mouse | Mouse NOD2 | No | No |
| PA5-19991 | Human | Synthetic peptide corresponding to 16 amino acids at the amino terminus of human NOD2 | No | No |
| PA5-19992 | Human | Synthetic peptide corresponding to 14 amino acids at the carboxy terminus of human NOD2 | No | No |
| MA1-16611 | Human, Mouse | Recombinant human NOD2 protein corresponding to residues 28-301 | No | No |
| PA5-111394 | Human | Recombinant protein corresponding to human NOD2 | No | No |
| H00064127-D01P | Human | NOD2 (AAI56572.1, amino acids ~1-1040) full-length human protein | No | No |
| NOD2-201AP | Mouse, Rat | Synthetic peptide taken within amino acid region 700-780 on mouse Nucleotide-binding oligomerization domain-containing protein 2 | No | No |
| PA5-18572 | Human | Synthetic peptide sequence (KFRFTDRERHCSPTD) corresponding to the internal amino acids of CARD15 | No | No |
| BS-7084R | Human, Mouse, Rat | KLH conjugated synthetic peptide derived from human CARD15, amino acids 251-350 | Yes | No |
| PA5-23101 | Human | Synthetic peptide corresponding to amino acids 125-140 of human NOD2 | No | No |

Across all antibodies tested, only a subset detected the intended human protein with specificity, and cross-reactivity with the zebrafish orthologs was rare. Of eight commercial antibodies tested against NOD1, five detected the human protein with specificity but none detected zebrafish NOD1. Among eleven commercial antibodies tested against NOD2, two detected their expected human protein target, whereas none detected the zebrafish ortholog. For NLRP1, three of the seven antibodies detected the human protein, and one antibody detected the zebrafish protein. Similarly, one of three antibodies raised against human NLRP14 detected both the human protein and the zebrafish protein si:ch73-233m11.2. Because available antibodies capable of recognizing the zebrafish orthologs of NLRP1 and si:ch73-233m11.2 were identified, these targets were deprioritized in subsequent aptamer development efforts. These results highlight the limited availability of reliable antibodies for detecting zebrafish NLR proteins and motivated the development of alternative affinity reagents.

### Identification of zebrafish NLR-binding aptamers using the APT-SNAP platform

To identify aptamers capable of recognizing zebrafish NLR proteins, we developed the APT-SNAP platform, a novel high-density DNA microarray designed for parallel screening of protein-aptamer interactions. The array contains approximately one million unique DNA features, wherein 10^5^-10^6^ replicate DNAs are synthesized within each circular 30 µm feature. Every feature or “address” on the array can be identified by a unique x,y coordinate. Probes within the features consist of two regions, a 40 nucleotide (nt) aptamer, whose sequence varies for each probe and folds into a unique secondary structure, and a 20-nt conserved spacer to elevate the probe from the array surface. Recombinant His-tagged NLR proteins were incubated on the arrays, and bound proteins were detected using a fluorescent anti-His-tag antibody (Fig. 3A).

**Figure 3.**
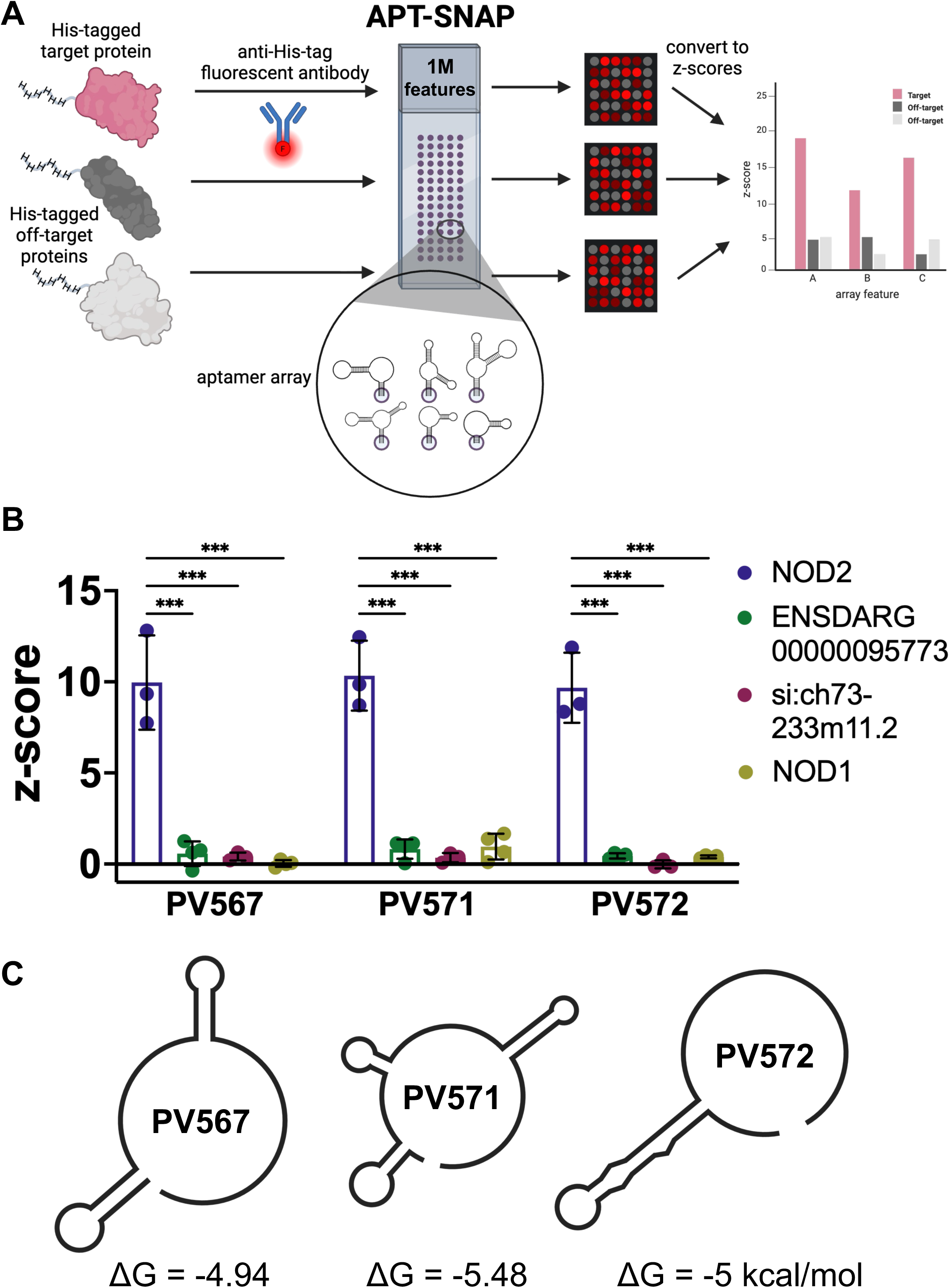
APT-SNAP screening identifies candidate NOD2-binding aptamers. A) Schematic representation of the APT-SNAP screening platform. His-tagged target or off-target proteins are incubated on high-density aptamer arrays containing approximately one million unique features. Protein binding is detected using fluorescent anti-His antibody and converted to z-scores to identify candidate aptamers. B) Z-scores from APT-SNAP screening for three candidate NOD2-binding aptamers (PV567, PV571, and PV572) tested against NOD2 and three off-target proteins (ENSDARG00000095773, si:ch73-233m11.2, and NOD1). Data represent replicate array experiments (n = 3-4). C) Predicted secondary structures and calculated free energy (ΔG) values for candidate aptamers determined using mFold.

APT-SNAP screening experiments were performed in triplicate for each NLR protein. Fluorescence intensity at each array feature was quantified and converted to z-scores to identify aptamers exhibiting selective binding to the target protein relative to counter-selection proteins (Fig. 3B, S1). Candidate aptamers were prioritized based on strong signal enrichment for the target protein compared with off-target NLR proteins.

Among the proteins screened, NOD2 produced the strongest and most reproducible binding signals across replicate experiments. Three DNA aptamers, designated PV567, PV571, and PV572, exhibited approximately five-fold higher signal intensity for NOD2 compared with the counter-selection proteins ENSDARG00000095773, si:ch73-233m11.2, and NOD1 (Fig. 3B, S1). Predicted secondary structures for each candidate aptamer were generated using mFold (29), revealing distinct stem-loops with calculated free energy values consistent with stable folding (Fig. 3C). Based on these results, subsequent validation experiments focused on NOD2 and three related NLR proteins (ENSDARG00000095773, si:ch73-233m11.2, and NOD1).

### Biochemical validation of candidate NOD2-binding aptamers by ELASA

Binding of candidate aptamers PV567, PV571, and PV572 to zebrafish NOD2 was first verified using enzyme-linked apta-sorbent assay (ELASA), an aptamer-based analogue of ELISA in which immobilized aptamers capture target proteins. In this assay format, biotinylated aptamers were immobilized on streptavidin-coated plates and incubated with purified His-tagged NLR proteins. Bound protein was detected using an anti-His-tag antibody followed by colorimetric detection.

All three candidate aptamers produced statistically significant signal when incubated with purified NOD2 compared with a biotinylated control DNA oligonucleotide containing a c-Myc-aptameric-binding motif (PV535) that matched in length to the candidate aptamers (Fig. 4 and Table S5). The candidate aptamers did not show detectable binding when incubated with two NLR family members (ENSDARG00000095773 and si:ch73-233m11.2 which bear the NACHT and LRR domains similar to NOD2) and the unrelated protein maltose binding protein (MBP). These results indicate that the candidate aptamers demonstrate specificity for NOD2 relative to related NLR proteins (Fig. 4). Representative experiments further demonstrated that aptamer binding to NOD2 is concentration-dependent, while maintaining little to no binding to the counter-selection proteins and MBP (Fig. S2).

**Figure 4.**
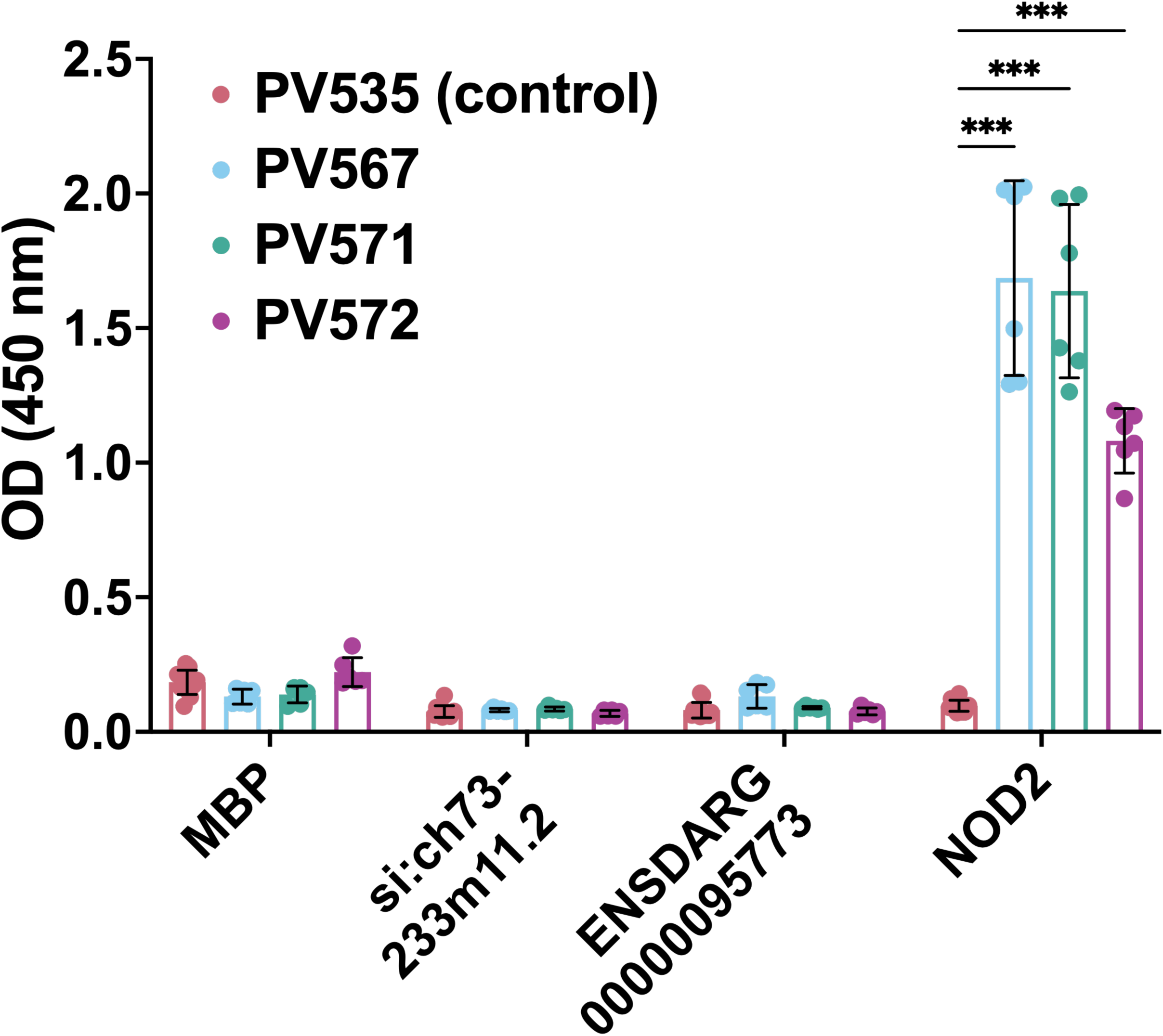
NOD2 aptamer candidates demonstrate target binding and specificity in ELASA format. Binding of candidate aptamers to purified proteins was measured using an enzyme-linked apta-sorbent assay (ELASA). Data represent mean ± SD from at least six technical replicates derived from a minimum of two independent experiments. Statistical comparisons were performed using two-tailed t-tests (n = 6; ***p < 0.001). Mean differences and p-values are reported in Table S5.

### Aptamer-mediated pull-down of NOD2

To determine whether the aptamers could capture NOD2 from complex mixtures, aptamer-mediated pull-down assays were performed using recombinant baculovirus infected insect cell lysates expressing recombinant NLR proteins. Biotinylated aptamers were immobilized on streptavidin magnetic beads and incubated with insect cell lysates. Following washing, bound proteins were eluted and analyzed by SDS-PAGE. All three candidate aptamers enriched NOD2 from recombinant lysates, whereas minimal enrichment was observed when lysates expressing counter-selection proteins ENSDARG00000095773 or si:ch73-233m11.2 were used (Fig. 5). These results demonstrate that the candidate aptamers can selectively capture NOD2 from protein lysates.

**Figure 5.**
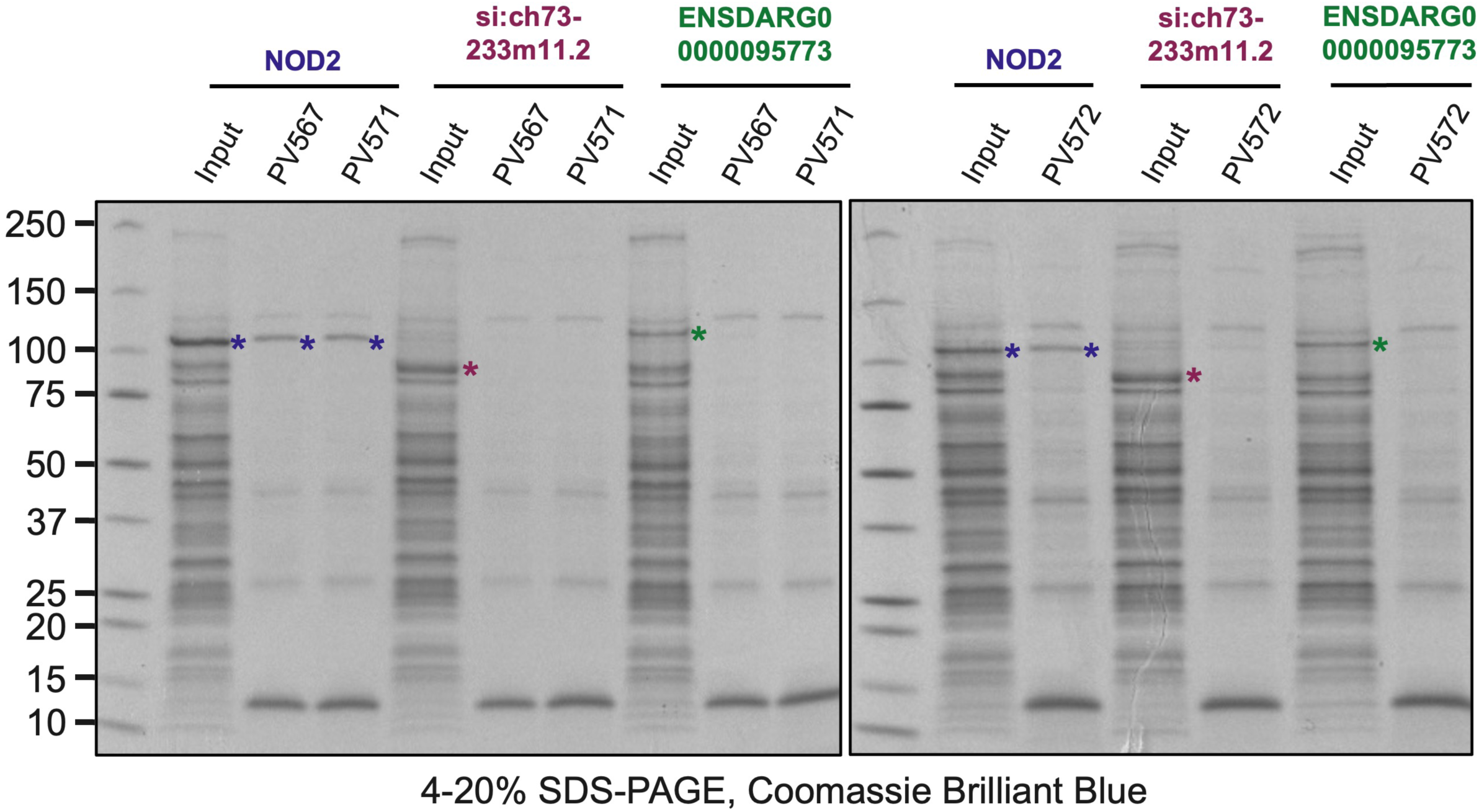
Pull-down assays demonstrate selective enrichment of NOD2 by candidate aptamers. Biotinylated aptamers were used as bait on streptavidin beads to capture proteins from insect cell lysates overexpressing NOD2 or off-target NLR proteins. Bound proteins were eluted from the beads and analyzed by SDS-PAGE followed by GelCode™ Blue Coomassie staining. Input lysates and aptamer-bound fractions are shown for NOD2 (indigo), si:ch73-233m11.2 (magenta), and ENSDARG00000095773 (green). Bands at the expected size for each protein are marked with an asterisk (*). NOD2 is enriched in the aptamer-bound fractions, whereas off-target proteins show no enrichment. The ∼13 kDa band corresponds to denatured streptavidin monomers from the beads.

### Quantitative binding analysis by biolayer interferometry

To quantify the binding affinity of the candidate aptamers for NOD2, we performed BLI. Biotinylated aptamers were immobilized on streptavidin biosensors and exposed to purified NOD2 at varying concentrations. Sensorgrams were fit using a global 1:1 binding model to determine kinetic parameters.

Aptamer PV567 exhibited high-affinity binding to NOD2 with an equilibrium dissociation constant (K_D_) of 431 pM, and PV571 bound with nanomolar affinity (K_D_ = 1.00 nM) (Fig. 6, S3, and Table 2). In contrast, PV535, the negative control biotinylated DNA, showed no measurable binding under identical conditions (Fig. S3). Although PV567 also bound to the related NLR protein ENSDARG00000095773 with an affinity (K_D_) of 2.21 nM (Fig. 6 and Table 2), the interaction was significantly weaker than with NOD2. Overall, PV567 interacted with NOD2 with a 5.13-fold higher affinity K_D_, a 3.07-fold faster association rate, and a 1.67-fold slower dissociation rate than when interacting with ENSDARG00000095773 (Table 2). Neither PV567 nor PV571 showed any measurable binding to NOD1 (Fig. 6, S3). These results indicate specific, high affinity binding of the candidate aptamers to NOD2.

**Figure 6.**
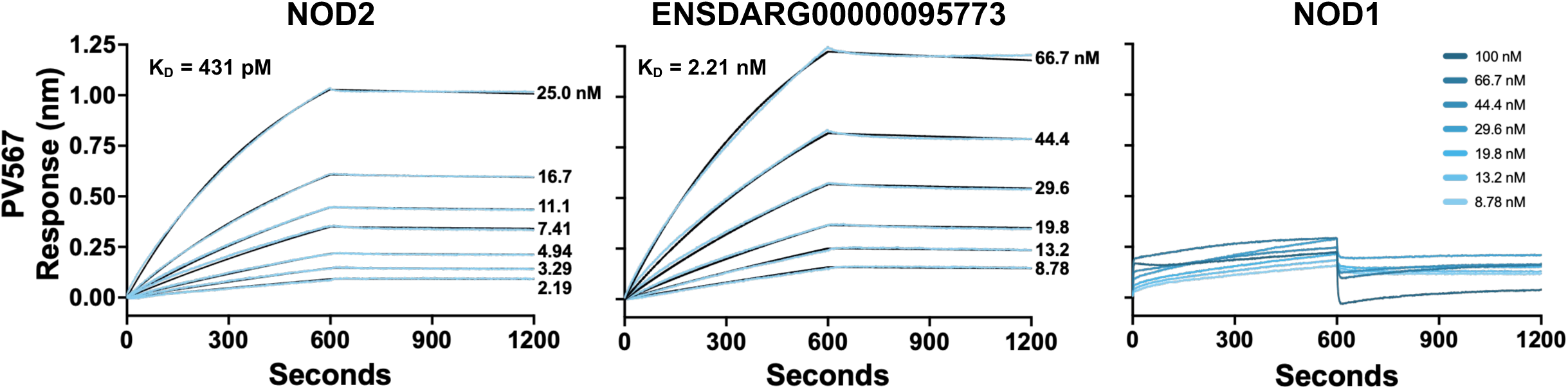
Biolayer interferometry analysis of aptamer binding. Streptavidin biosensors were loaded with biotinylated DNA aptamers and exposed to purified protein at the indicated concentration ranges. Background-subtracted sensorgrams and global 1:1 kinetics fits (black) are shown. Aptamer PV567 binding to NOD2 (K_D_ = 431 pM), off-target protein ENSDARG00000095773 (K_D_ = 2.21 nM), and no observable binding to off-target protein NOD1. Experiments were performed at 30 °C in Octet kinetics buffer supplemented with 5 mM MgCl_2_.

**Table 2.** Biolayer interferometry kinetic parameters for aptamer-protein interactions.

| **Aptamer** | **Protein** | **K<sub>D</sub>** | **k<sub>on</sub> (M<sup>-1</sup> s<sup>-1</sup>)** | **k<sub>off</sub> (s<sup>-1</sup>)** | **χ<sup>2</sup>** | **R<sup>2</sup>** |
|---|---|---|---|---|---|---|
| PV567 | NOD2 | 431 ± 11 pM | 8.29×10<sup>4</sup> | 3.58×10<sup>-5</sup> | 0.3032 | 0.9995 |
| PV571 | NOD2 | 1.00 ± 0.01 nM | 1.22×10<sup>5</sup> | 1.22×10<sup>-4</sup> | 1.042 | 0.9983 |
| PV567 | ENSDARG00000095773 | 2.21 ± 0.04 nM | 2.70×10<sup>4</sup> | 5.98×10<sup>-5</sup> | 0.6308 | 0.9993 |

### Aptamers recognize native but not denatured NOD2

The preceding assays were performed under native protein conditions, so the aptamers were also evaluated to determine whether they could detect NOD2 by western blot. Despite extensive testing, the aptamers did not produce detectable signal under denaturing SDS-PAGE conditions (data not shown). This outcome is consistent with the APT-SNAP screening format, which presents folded proteins and therefore allows for aptamers that recognize conformational epitopes. Together with the ELASA, pull-down, and BLI results, these findings indicate that the identified aptamers bind native NOD2 with high affinity and specificity but may not recognize denatured NOD2.

### Aptamer-based detection of zebrafish NOD2 in cellular assays

To determine whether the aptamers could detect NOD2 in a cellular context, we performed fluorescence microscopy using biotinylated PV567 followed by staining with Alexa Fluor™ 647-labeled streptavidin. ZF4 zebrafish cells, which are known to constitutively express NOD2 protein (30), were stained with the aptamers and a scrambled version of PV567 as a negative control (PV682). Representative images of PV567 and PV682 show aptamer-specific staining within the cells, using phalloidin to stain actin and DAPI to visualize the nuclei (Fig. 7A). Quantification of fluorescence intensity normalized to nuclear area showed that PV567 signal was significantly higher than that observed for the control oligonucleotide PV682 (Fig. 7B and S4). In contrast, PV571 and PV572 did not display any increase in fluorescence relative to PV682.

**Figure 7.**
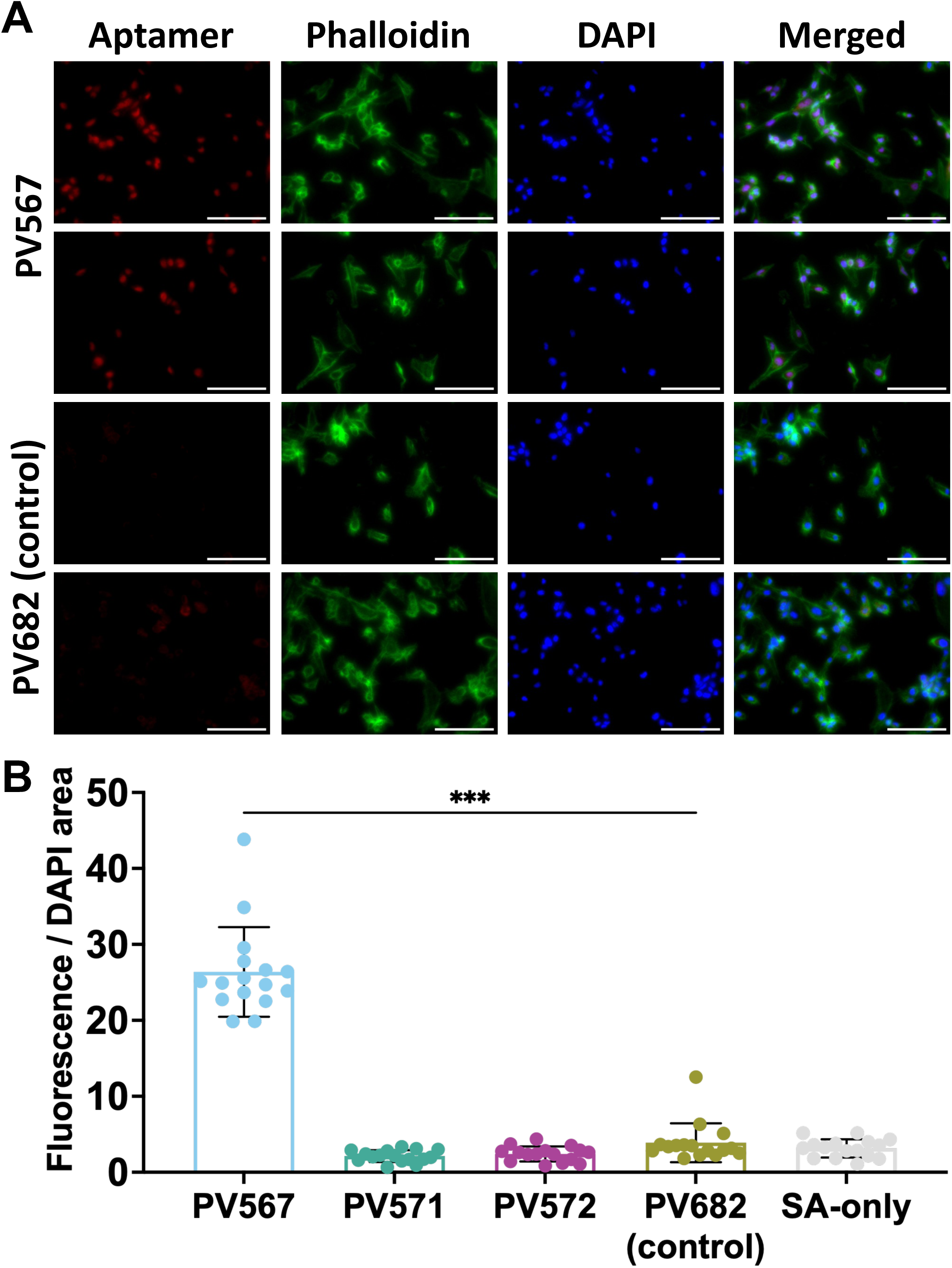
Fluorescence microscopy detection of NOD2 using candidate aptamers. A) Representative fluorescence microscopy images of cells stained with biotinylated aptamer (PV567) or negative control (PV682) followed by staining with streptavidin-Alexa Fluor™ 647 conjugate. Aptamer signal (red), actin cytoskeleton staining with phalloidin (green), and nuclear staining with DAPI (blue) are shown along with merged images. Images represent maximum contrast projections of 12 z-stack images acquired at 20 µm intervals. B) Quantification of fluorescence signal normalized to DAPI area for each aptamer condition. Data are presented as individual measurements with mean ± SD. Statistical significance was determined using t-tests (n = 16; ***p < 0.001).

## DISCUSSION

A major barrier to studying innate immune signaling pathways in zebrafish is the limited availability of reliable protein detection reagents. Although many NLRs are conserved between mammals and zebrafish, sequence divergence between orthologs can substantially reduce the effectiveness of antibodies developed against mammalian proteins for use in zebrafish. Consistent with this limitation, our evaluation of commercially available antibodies raised against human NLR proteins demonstrated very limited cross-reactivity with zebrafish orthologs. Among the antibodies tested, only 38% (11 of 29) detected their intended human targets with specificity despite using manufacturers’ recommended experimental conditions, and recognition of the corresponding zebrafish proteins was rare. These findings highlight the scarcity of validated detection reagents for zebrafish NLRs and underscore the need for alternative affinity reagents for studying innate immune pathways in this model system.

In this study, we established a workflow for the identification and validation of aptamer-based detection reagents targeting zebrafish NLR proteins. Using our novel APT-SNAP platform, a high-density aptamer microarray technology that enables parallel screening of protein interactions with one million user-defined aptamers wherein each aptamer is located at a specific x-y coordinate, we identified multiple candidate aptamers capable of binding purified zebrafish NLR proteins. Subsequent validation experiments focused on zebrafish NOD2, which produced robust and reproducible binding signals from screening on a single array design. Three candidate DNA aptamers were identified and further evaluated using orthogonal biochemical and cellular assays.

Biochemical validation demonstrated that the candidate aptamers specifically recognize zebrafish NOD2 under native conditions. In ELASA experiments, all three aptamers produced strong, concentration-dependent binding signals for NOD2 while showing minimal signal for related NLR proteins or an unrelated control protein (MBP). Aptamer-mediated pull-down assays further demonstrated that the candidate sequences were capable of enriching NOD2 from recombinant insect cell lysates, indicating that the aptamers can capture their target from complex protein mixtures. Quantitative analysis using BLI revealed high-affinity interactions between the aptamers and NOD2, with the strongest candidate exhibiting sub-nanomolar affinity. Although aptamer PV567 also displayed detectable binding to a closely related NLR protein, the interaction occurred with weaker affinity, indicating preferential recognition of NOD2 relative to related family members.

Notably, the aptamers identified in this study did not produce detectable signal in western blot assays performed under denaturing conditions. This observation is consistent with the screening format used during APT-SNAP array experiments, which presents folded proteins and therefore favors aptamers that recognize conformational epitopes. Together with the ELASA, pull-down, and BLI results, these findings indicate that the identified aptamers bind native NOD2 with high affinity and specificity but may not recognize the denatured form of the protein.

Aptamer PV567 was also capable of detecting NOD2 in a cellular context. Fluorescence microscopy experiments demonstrated that the lead aptamer produced strong staining consistent with the expected expression of NOD2 in cultured ZF4 embryonic fibroblast cells (30), while a scrambled control DNA did not. Our results suggest that NOD2 may be both cytoplasmic and nuclear in ZF4 cells. Previous studies have shown that NOD2 is primarily located in the cytoplasm (31, 32). However, other studies identified human NOD2 interaction with nuclear lamins in hepatocellular carcinoma samples and verified the presence of a functional nuclear localization signal (NLS) (33). Using a similar cNLS mapping tool (34), zebrafish NOD2 contains a bipartite NLS of PSLKRYLRKEVLLKGFSPGGIDCFVKKHYSD starting at position 410 in the protein. These findings support that zebrafish NOD2 may localize to both the cytoplasm and nucleus depending on cellular and experimental conditions, and this warrants further investigation in future studies. These results support that aptamer-based detection reagents identified through the APT-SNAP platform can function in both biochemical assays and cellular imaging applications.

Beyond the identification of NOD2-binding aptamers, this work establishes a generalizable workflow for the development of affinity reagents targeting proteins that lack reliable antibodies. Traditional aptamer discovery methods typically rely on iterative enrichment approaches such as Systematic Evolution of Ligands by EXponential enrichment (SELEX), which can require multiple rounds of selection and screening (35–37). In contrast, the APT-SNAP platform enables direct, high-throughput interrogation of large libraries of folded nucleic acid structures, which facilitates rapid identification of candidate aptamers without iterative enrichment. Of note, this APT-SNAP workflow successfully identified aptamers by screening just 10^6^ sequences compared with a starting pool of 10^14^ sequences selected from in SELEX. In addition, the purified zebrafish NLRs that served as targets or counter-selection proteins provide another resource for the zebrafish research community.

Additional work will be required to evaluate the performance of these aptamers in detecting endogenous NOD2 in native biological contexts beyond the purified proteins, overexpression-based systems, and ZF4 cells described here. Future studies could also explore additional rounds of array-based screening or a combination of APT-SNAP with enrichment-based selections such as SELEX to maximize the advantages of both methods for further optimization of the specificity of candidate aptamers. While careful monitoring of aptamer selections or the use of modified nucleic acids can increase the rates of successful SELEX, the often-cited success rate is only 30% (38–40). The APT-SNAP approach either alone or in combination with reduced rounds of SELEX may improve aptamer identification success rates without the need to use expensive modified nucleic acids.

In conclusion, this study establishes a workflow for the identification and validation of DNA aptamers, demonstrated here using zebrafish NLR proteins. The aptamers identified in this study provide new molecular tools for studying NOD2 in the zebrafish model system. This work positions APT-SNAP as a complementary method to other aptamer discovery platforms like SELEX by enabling direct screening of structurally defined DNA probes against purified native proteins, followed by rapid validation in biochemical and cellular assays. More broadly, the strategy described here may be extended to other challenging protein targets that lack reliable antibody reagents, expanding the toolkit available for studying protein function in diverse and underserved biological systems.

## EXPERIMENTAL PROCEDURES

### Reagents

General biological reagents and chemical compounds were purchased from commercial suppliers including Thermo Fisher Scientific, Promega, Sigma-Aldrich/Merck, New England Biolabs, and Bio-Rad Laboratories.

The following primary antibodies were used: FLAG (Origene, TA592569; Cell Signaling Technology, 2368S), HRP-conjugated PentaHis (Qiagen, 34460), and Alexa Fluor™ 647-conjugated PentaHis (Qiagen, 35370). HRP-conjugated secondary antibodies included anti-mouse IgG (Promega, W402B; 1:2500), anti-rabbit IgG (Promega, W401B; 1:2500), goat anti-rat IgG (Thermo Fisher Scientific, 31470; 1:10,000), and rabbit anti-goat IgG (Thermo Fisher Scientific, 31402; 1:10,000). Mammalian NLR antibodies tested in western blot experiments were purchased from Thermo Fisher Scientific. Catalog numbers and additional details for these antibodies are provided in Tables 1, S2, S3, and S4. Antibodies were diluted according to the manufacturers’ instructions unless otherwise specified above.

DNA aptamers were synthesized as single-stranded oligonucleotides by Integrated DNA Technologies (IDT), including versions containing modifications such as 3’ biotin or fluorescein labels when required for downstream assays.

### Protein cloning and expression using BEVS

Expression of His-tagged recombinant proteins was performed using the ExpiSf Expression System (Gibco, Thermo Fisher Scientific) and the Bac-to-Bac Baculovirus Expression System (Invitrogen, Thermo Fisher Scientific) according to the manufacturers’ instructions. The protein coding sequence for each of the zebrafish target and counter-selection genes were obtained from the UniProt database (accession numbers listed in Table S1). The corresponding nucleic acids sequence was codon-optimized for expression in *Spodoptera frugiperda* 9 (Sf9) cells. They were then synthesized as ∼700 bp eBlocks (IDT) and cloned into the pFastBac HT expression vector (Invitrogen, Thermo Fisher Scientific) using the NEBuilder HiFi DNA Assembly kit (New England Biolabs). Constructs were transformed into competent *E. coli*, and plasmid DNA was prepared for bacmid generation. Bacmid DNA containing the integrated gene of interest was transfected into ExpiSf9 insect cells (Thermo Fisher Scientific, A35243) to generate recombinant baculovirus.

Viral stocks were amplified through two to three passages to obtain high-titer virus, which was quantified using the BacPAK Baculovirus Rapid Titer Kit (Takara Bio USA). Expression tests were performed in ExpiSf9 and High Five™ (Hi5) insect cells (Thermo Fisher Scientific, B85502). Multiplicity of infection (MOI; 1, 5, and 10) and harvest time (24, 48, and 72 hours post-infection) were evaluated to optimize expression conditions.

Protein expression levels were assessed by western blot analysis, and protein integrity was evaluated by SDS-PAGE with Coomassie staining. Optimal infection conditions were used to generate larger-scale cultures for protein purification and downstream experiments.

### Protein purification

Baculovirus-infected cells were resuspended in Lysis Buffer 1 (PBS, 0.5% NP-40, and 1 mM imidazole, 10 mM KCl, 1 mM MgCl_2_, 0.1 mM ADP, and 0.1 mM DTT) supplemented with Pierce protease inhibitor tablets (Thermo Fisher Scientific). After incubation on ice for 5 min, lysates were clarified by centrifugation at 20,000 × g for 5 min at 4 °C, and the supernatant containing the soluble protein fraction was collected. The clarification step was repeated with a 10-min centrifugation to complete removal of cellular debris. The supernatant was applied to Ni Sepharose resin (Cytiva) pre-equilibrated with lysis buffer and centrifuged through a small microcentrifuge column cup at 50-500 × g as needed for flow-through. The column was washed with Wash Buffer 1 (300 mM NaCl, 0.01% Tween-20, 10 mM KCl, 1 mM MgCl_2_, 0.1 mM ADP, 0.1 mM DTT, and 1 mM PMSF) containing 10-50 mM imidazole. Bound proteins were eluted using Wash Buffer 1 containing 200 mM imidazole. Eluted proteins were buffer exchanged into Storage Buffer (20 mM HEPES, 150 mM NaCl, 10 mM KCl, 1 mM MgCl_2_, 0.1 mM ADP, 0.1 mM DTT, 0.01% Tween-20, and 20% glycerol). Purified proteins were aliquoted, flash frozen in liquid nitrogen, and stored at -80 °C.

Protein purity and quality were evaluated by SDS-PAGE using GelCode™ Blue Stain Reagent (Thermo Fisher Scientific), a high sensitivity Coomassie G-250-based stain, and by western blot analysis as described below. Because purity levels varied between NLR targets and counter-selection proteins, protein concentrations were estimated by SDS-PAGE densitometry relative to BSA standards prior to use in validation assays.

### Western blot analysis

Western blot analysis of overexpression lysates and purified zebrafish NLR proteins was performed using the PentaHis HRP Conjugate Kit (Qiagen, 34460) according to the manufacturer’s instructions. Commercially available antibodies raised against mammalian NLR proteins (Thermo Fisher Scientific) were evaluated for cross-reactivity with zebrafish orthologs. A panel of 29 antibodies targeting human NOD1, NOD2, NLRP1, and NLRP14 (the human homolog of zebrafish si:ch73-233m11.2) was tested. Notably, 26 of the 29 antibodies were listed by the manufacturer as validated for western blot applications.

FLAG-tagged overexpression lysates of human NOD1, NOD2, NLRP1, and NLRP14 were obtained from Origene and tested alongside His-tagged overexpression lysates of zebrafish NOD1, NOD2, NLRP1 and si:ch73-233m11.2 generated in-house. Zebrafish lysates were prepared by suspending cell pellets in RIPA buffer supplemented with Pierce nuclease and protease/phosphatase inhibitor cocktail (Thermo Fisher Scientific). Samples were incubated at room temperature for 10 min followed by an additional 30 min on ice. Lysates were clarified by centrifugation at 14,000 × g for 10 min, and protein concentrations were determined using a BCA assay.

Lysate samples (5-20 µg depending on expression level) were separated by SDS-PAGE using precast 4-20% gradient gels (BioRad) and transferred to PVDF membranes using the Trans-Blot Turbo transfer system (BioRad), with transfer efficiency confirmed by Ponceau S staining. Membranes were washed with TBST and blocked in 5% non-fat milk in TBST for 1 hour at room temperature with gentle rocking. After blocking, membranes were incubated overnight at 4 °C with primary antibodies diluted in blocking buffer. The following day, membranes were equilibrated to room temperature, washed with TBST, and incubated with appropriate HRP-conjugated secondary antibodies for 1 hour. After additional washes with TBST, protein bands were visualized using Clarity Western ECL substrate (Bio-Rad) and imaged using a ChemiDoc™ MP imaging system (Bio-Rad) with Image Lab software.

### APT-SNAP microarray design

A custom DNA microarray was designed for initial aptamer screening (41). Each array contained approximately one million features, with each feature composed of ∼10^5^-10^6^ identical 60-nucleotide (nt) DNA probes immobilized via their 3’ terminus. To design the array and generate candidate aptamer sequences, 100 million random 40-nt DNA sequences were computationally generated and analyzed using the UNAFold software suite in batches of one million sequences. Sequences predicted to adopt a single dominant secondary structure were selected to ensure structural homogeneity across the array library. Selected sequences were classified according to their predicted secondary structures based on Vienna notation generated by UNAFold. Structural classes included stem-loops (hairpins), bubbles and bulges of variable lengths and sequences, and three-way and four-way junctions (cruciform) of variable arm lengths and sequences. In addition, intramolecular G-quadruplex and i-motif structures were incorporated, including designs containing two to eight tetrads with variable linker lengths and sequences between strands. Each probe consisted of a 40-nt variable aptamer region preceded by a constant 20-nt spacer sequence positioned proximal to the array surface to elevate the aptamer structure away from the surface and improve accessibility for protein binding. Custom arrays based on this design were synthesized by Agilent Technologies.

### APT-SNAP microarray binding

To induce aptamer folding, arrays were first incubated in preheated 7M Urea in 1× PBS at 65 °C for 30 min with brief agitation at 10-min intervals. Arrays were then transferred to preheated 1× PBS and incubated at 65 °C for an additional 15 min with brief agitation halfway through the incubation. Arrays were subsequently incubated in Non-Stringent Wash Buffer (6× SSPE, 0.01% Tween-20 in ultrapure water) for 15 min at room temperature protected from light. Arrays were then briefly rinsed in ice-cold Final Wash Buffer (FWB; 0.6× SSC) and dried. A hybridization chamber was placed over the array surface and hydrated twice with ultrapure water before blocking with 2.5% fat-free milk for 1 hour with rotation. After blocking, the hybridization chambers were rinsed with Binding Buffer (50 mM HEPES, 100 mM KCl, 0.1 mM EDTA, 5 mM MgCl_2_, 0.0035% NP-40, and 0.25 mM DTT). Purified zebrafish NLR protein was diluted in ice-cold Binding Buffer (NOD2: 4 nM; si:ch73-233m11.2: 10 nM; ENSDARG00000095773: 10 nM) together with a 1:833 dilution of Alexa Fluor™ 647-conjugated PentaHis antibody (Qiagen) and incubated on the arrays for 1 hour with rotation. After incubation, the protein mixture was removed, and the hybridization chambers were rinsed with ice-cold Binding Buffer. The hybridization chambers were then removed from the arrays, and the arrays were briefly rinsed in ice-cold FWB, dried, and scanned using a GenePix® 4000B scanner (Molecular Devices) with GenePix® Pro 7 image acquisition software (Molecular Devices), using our established array procedures (42).

Array data were extracted using Agilent Feature Extraction Software (version 12.1). For each array, a z-score was calculated for every feature as z = (x − μ) / σ, where x is the feature’s median fluorescence signal, and μ and σ are the mean and standard deviation of median signal across all features on that array. Replicate arrays were scored independently before comparison across proteins.

### Enzyme-Linked Apta-Sorbent Assay (ELASA)

Biotinylated aptamers (IDT) were diluted in 0.1× TE buffer and heated at 95 °C for 5 min to denature secondary structures. Four volumes of ice-cold aptamer Folding Buffer (62.5 mM HEPES, 125 mM KCl, 0.125 mM EDTA, and 6.25 mM MgCl_2_) were then added, and samples were incubated on ice for 5 min to allow refolding. Folded aptamers were applied to streptavidin (SA)-coated 96-well plates (Thermo Fisher Scientific, 15121) to a final concentration of 100 nM (200 µL per well) and incubated for 30 min at room temperature. Wells were washed twice with Wash Buffer 2 (50 mM HEPES, 100 mM KCl, 0.1 mM EDTA, 5 mM MgCl_2_, and 0.0035% NP-40; 300 µL per well) and then blocked with 2.5% fat-free milk in wash buffer (200 µL per well) for 30 min at room temperature. During the blocking step, HRP-conjugated PentaHis antibody (Qiagen; 1:2000 dilution) was pre-incubated with purified NLR protein (2.5 nM) in ice-cold ELASA Buffer (Wash Buffer 2 supplemented with 10 µg/mL salmon sperm DNA and 1 mg/mL BSA) for approximately 10 minutes on a rotator. After blocking, plates were washed three times with Wash Buffer 2 (300 µL per well), and 100 µL of the protein-antibody mixtures were added to appropriate wells. Plates were incubated for 30 min at room temperature and then washed three times with Wash Buffer 2 (300 µL per well). TMB Substrate (Thermo Fisher Scientific; 100 µL per well) was added and allowed to develop for 10 min. The reaction was stopped by the addition of 100 µL of 1 M HCl, and the absorbance at 450 nm was measured using a Tecan Infinite® M1000 Pro plate reader with Magellan™ software (Tecan).

### Pull-down assay using aptamers

Streptavidin magnetic beads (Promega, Z5481) were washed three times with 0.5× SSC and once with Wash Buffer 3 (50 mM HEPES, 100 mM KCl, 0.1 mM EDTA, 5 mM MgCl_2_, 0.0035% NP-40, and 1 mM DTT) before being resuspended to their original volume. Biotinylated aptamers were refolded as described in the ELASA section prior to use. Cell pellets were resuspended in Lysis Buffer 2 (20 mM HEPES, 50 mM NaCl, 0.07% NP-40, and 1× protease inhibitor) and incubated on ice for 5 min before clarification by centrifugation at 14,000 × g for 5 min at 4 °C. The supernatant was collected and protein concentration was estimated by absorbance at 280 nm prior to use in pull-down experiments. Aptamer in excess of the bead binding capacity was incubated with 20 µg of streptavidin beads in 100 µL of Wash Buffer 3 for 15 min on a rotator to allow immobilization. Beads were then collected using a magnetic stand, and unbound aptamer was discarded. For each NLR target, 40 µg of baculovirus overexpression lysate was diluted in 0.5 mL of Wash Buffer 3 supplemented with 1× protease inhibitor and incubated with aptamer-coated beads for 1 hour with rotation. Beads were subsequently collected using a magnetic stand and washed three times with 200 µL of wash buffer. Bound proteins were eluted by incubating beads in 30 µL of reducing Laemmli sample buffer (50 mM DTT) at 95 °C for 2 min. Input lysates and eluates were analyzed by SDS-PAGE with GelCode™ Blue Coomassie stain.

### Bio-layer Interferometry (BLI)

Kinetic binding measurements were performed using an Octet® Red96 instrument (Sartorius/ForteBio) with streptavidin biosensors (Sartorius, 18-5019). DNA aptamers containing a 3’ biotin modification were immobilized onto streptavidin biosensors following heat-refolding. Aptamers were diluted to 500 nM in 1× Octet® Kinetics Buffer (Sartorius, 18-1105) supplemented with 5 mM MgCl_2_, heated at 90 °C for 10 min, and snap-cooled on ice prior to immobilization. Purified recombinant zebrafish NOD2 protein (prepared as described above) was used as the analyte and diluted into the same Mg^2+^-supplemented kinetics buffer. All BLI experiments were conducted at 30 °C with orbital shaking at 1000 rpm. The assay consisted of the following steps: baseline (180 s), aptamer loading (120 s), baseline (180 s), association (600 s), and dissociation (600 s). Background subtraction was performed using buffer-only reference wells.

Processed sensorgrams were analyzed using Octet® Data Analysis Software (version 12). Association and dissociation curves were globally fit using a 1:1 binding model. Reported kinetic parameters include the association rate constant (K_on_), dissociation rate constant (K_off_), and equilibrium dissociation constant (K_D_ = K_off_/K_on_). Nonspecific binding was assessed using the c-Myc negative control aptamer described in the Results section.

### Fluorescence microscopy

ZF4 zebrafish fibroblast cells (ATCC, CRL-2050) were cultured according to the supplier’s recommendations in DMEM:F12 (Gibco, 11320033) supplemented with 10% FBS (Gibco, A5209401) and 1% Pen/Strep (Gibco, 15140122) at 5% CO_2_ and 28 °C. Cells were seeded in a 24-well plate 48 hours prior to fixation. Cells were rinsed with PBS and fixed in 3.7% paraformaldehyde (Thermo Fisher Scientific, J61984.AK) for 10 min at room temperature. Cells were then permeabilized with 0.1% Triton X-100 in PBS for 5 to 10 min, followed by one wash with PBS.

Biotinylated aptamers were heat-refolded prior to use as described above. Refolded aptamers were diluted to 100 nM in a solution containing 0.1 mg/mL BSA and 0.1 mg/mL yeast tRNA to reduce nonspecific binding. Cells were incubated with aptamers for 1.5 hours at room temperature. Scrambled aptamer and no-aptamer (streptavidin only) negative controls were included. Following aptamer incubation, cells were washed with PBS containing 5 mM MgCl_2_ and incubated with streptavidin Alexa Fluor™ 647 conjugate (Thermo Fisher Scientific, S21374) diluted 1:500 in PBS for 30 min at room temperature protected from light. Cells were then washed with PBS and stained with DAPI (Millipore Sigma, D9542; 1:10,000 dilution in PBS) for 10 min prior to final PBS washes. Cells were maintained in PBS for imaging. Another set of wells were treated as described above with additional staining with ActinGreen 488 ReadyProbe™ solution (Invitrogen, R37110). One drop per mL of PBS was mixed, added to cells, and incubated at room temperature for 30 min protected from light. Cells were washed again with PBS and maintained in PBS for imaging.

Fluorescence images were acquired using a BZ-X1000 fluorescence microscope (Keyence) equipped with a BZ-X Plan Apochromat 10× and Plan Fluorite 40× LD PH objectives (BZ-PA10; BZ-PF40LP) and a multi-band fluorescence filter set compatible with DAPI, Alexa Fluor™ 647, and ActinGreen 488 ReadyProbe™ (Multi-band Type 89000, OP-89167; HR filter GFP, OP-89162). For each well, the microscope automatically sampled 16 image fields and collected Z-stack images consisting of 12 optical sections spaced 20 µm apart using the 10× objective. Additional representative fields were captured using the 40× objective. Final images were generated using maximum contrast projection of the Z-stacks.

Fluorescence quantification was performed using the Image Cytometry module with the Hybrid Cell Count application (Keyence). Total Alexa Fluor™ 647 fluorescence and DAPI-positive nuclear area were measured for each image field. Signal intensity was normalized by calculating the ratio of total Alexa Fluor™ 647 fluorescence to total DAPI area for each field prior to statistical analysis. Representative images were processed using FIJI (ImageJ) for pseudocoloring, channel merging, and brightness/contrast adjustment. These adjustments were applied uniformly across each channel and did not affect the quantitative analysis.

## DATA AVAILABILITY

All data supporting the statements and conclusions made by the authors are included in the figures and in the supplemental information.

## SUPPORTING INFORMATION

This article contains supporting information.

## ACKNOWLEDGMENTS

The authors thank Keyence scientist, Conor Schlichting, for assistance with the microscopy studies and the staff at the University of Wisconsin-Madison Biophysics Instrumentation Facility for assistance with the BLI instrument.

## AUTHOR CONTRIBUTIONS

Conceptualization: CLW, MSO. Funding acquisition: CLW, MSO. Investigation: PEM, KSP, DTV, BJC, VK, ESO. Visualization: PEM, KSP. Data analysis: PEM, KSP, DTV, BJC, NRN, VK, ESO. Writing - original draft: PEM, KSP. Writing - review & editing: PEM, KSP, CLW, MSO, DTV, BJC, NRN, VK, ESO.

## FUNDING AND ADDITIONAL INFORMATION

This work was supported by a National Institutes of Health contract (75N93021C00055-0-9999-1) to MSO. The content is solely the responsibility of the authors and does not necessarily represent the official view of the NIH. BLI data were obtained at the University of Wisconsin-Madison Biophysics Instrumentation Facility, which was established with support from the University of Wisconsin-Madison and grants BIR-9512577 (NSF) and S10RR013790 (NIH).

