## Supporting Information for "Identification of Functional NOD2-Specific Aptamers Using APT-SNAP Microarrays to Screen Zebrafish Nod-like Receptors"

**Table S1. Summary of recombinant proteins.**

| Protein | Uniprot Accession Number | Predicted molecular weight (kDa) | Cell Line | Quantity Loaded for Western Blot (ng) |
|---|---|---|---|---|
| NOD1 (Origene) | A0A8M1RMD1 | 105 | – | 100 |
| NOD1 | X1WGQ4* | 105 | Hi5 | 100 |
| NOD2 | F8W3K2 | 112 | Sf9 | 100 |
| NLRP1 | A0A386CAB9 | 157 | Sf9 | 120 |
| si:ch73-233m11.2 | F1QZJ8 | 101 | Sf9 | 100 |
| si:ch211-214c20.1 | E9QC36* | 118 | Sf9 | 75 |
| ENSDARG00000095773 | B3DK63 | 123 | Hi5 | 10 |
| NLRC3L1 | E7FBD8 | 90 | Hi5 | 5 |
| NLRC6 | A5PF24 | 108 | Sf9 | 75 |
| si:ch211-195h23.3 | B0V1H3 | 57 | Hi5 | 10 |

*Entry was later removed from UniProt and replaced by A0A8M1RMD1

**Table S2. Human NOD1 antibody testing summary.**

| Antibody | Species Reactivity | Immunogen | Detects Human NOD1? | Detects Zebrafish NOD1? |
|---|---|---|---|---|
| PA5-79747 | Human, Rat | *E. coli*-derived human CARD4 recombinant protein (Position: M1-M160) | Yes | No |
| PA1-41036 | Human | Synthetic peptide corresponding to residues 930-949 of human NOD1 protein | No | No |
| PA5-21981 | Human | Recombinant fragment corresponding to a region within amino acids 698 and 953 of NOD1 | Yes | No |
| PA5-21015 | Human, Mouse, Rat | Synthetic peptide corresponding to 16 amino acids near C-terminus of human NOD1 | Yes | No |
| PA5-17328 | Human, Mouse, Non-human primate, Rat | Synthetic peptide corresponding to residues near the carboxy terminus of NOD1 protein | Yes | No |
| PA5-18027 | Human | Synthetic peptide sequence (EAKVYEDEKRIICF) corresponding to the C-terminus amino acids of NOD1 | No | No |
| BS-7085R | Human, Mouse, Rat | KLH conjugated synthetic peptide derived from human CARD4, amino acids 51-150 | Yes | No |
| H00010392-B01P | Human | NOD1 (AAH40339.1, amino acids ~1-953) full-length human protein | No | No |

**Table S3. Human NLRP1 antibody testing summary.**

| Antibody | Species Reactivity | Immunogen | Detects Human NLRP1? | Detects Zebrafish NLRP1? |
|---|---|---|---|---|
| PA5-116672 | Human, Mouse, Rat | Recombinant fusion protein corresponding to amino acids 1370-1473 of human NALP1 | Yes | Yes |
| 12256-1-AP | Human, Mouse | NLRP1 Fusion Protein Ag2908 (amino acids 1-146 encoded by BC016965) | No | No |
| PA5-20005 | Human | Peptide corresponding to amino acids 1461-1473 of human NLRP1 | No | No |
| PA5-111579 | Human | Recombinant protein corresponding to Human NLRP1, amino acids 1130-1261 | Yes | No |
| PA5-100640 | Human | A synthesized peptide derived from human NLRP1, corresponding to a region within N-terminal amino acids | No | No |
| 24792-1-AP | Human | NLRP1 Fusion Protein Ag20516 (amino acids 1031-1375 encoded by BC051787) | Yes | No |
| BS-6854R | Human, Mouse, Rat | KLH conjugated synthetic peptide derived from human NLRP1, amino acids 301-400 | No | No |

**Table S4. Human NLRP14 antibody testing summary.**

| Antibody | Species Reactivity | Immunogen | Detects Human NLR (NLRP14)?* | Detects Zebrafish NLR (si:ch73-233m11.2)? |
|---|---|---|---|---|
| PA5-21029 | Human, Mouse, Rat | A 16 amino acid peptide near the amino terminus of human NLRP14 | No | No |
| PA5-58802 | Human | Recombinant protein corresponding to Human NLRP14 | No | No |
| BS-19286R | Human, Mouse | KLH conjugated synthetic peptide derived from human NLRP14, amino acids 81-180 | Yes | Yes |

*Results of testing performed using human NLRP14-overexpressing lysate purchased from Origene. The positive control using the FLAG-tag antibody was inconclusive.

**Table S5. Additional statistical information for figure 4 ELASA.**

| Comparison | Protein | Mean difference | t-test p-value | Summary |
|---|---|---|---|---|
| PV567 vs PV535 (control) | MBP | 0.0409 | 0.1353 | ns |
| | si:ch73-233m11.2 | 0.0059 | 0.6091 | ns |
| | ENSDARG00000095773 | -0.03738333 | 0.08396 | ns |
| | NOD2 | -1.574267 | 0.0001058 | *** |
| PV571 vs PV535 (control) | MBP | 0.03318333 | 0.07724 | ns |
| | si:ch73-233m11.2 | 0.001616667 | 0.8921 | ns |
| | ENSDARG00000095773 | 0.004416667 | 0.7624 | ns |
| | NOD2 | -1.525617 | 0.0000696 | *** |
| PV572 vs PV535 (control) | MBP | -0.02578333 | 0.3833 | ns |
| | si:ch73-233m11.2 | -0.003783333 | 0.4748 | ns |
| | ENSDARG00000095773 | -0.00915 | 0.07032 | ns |
| | NOD2 | -0.9985333 | 0.00000419 | *** |

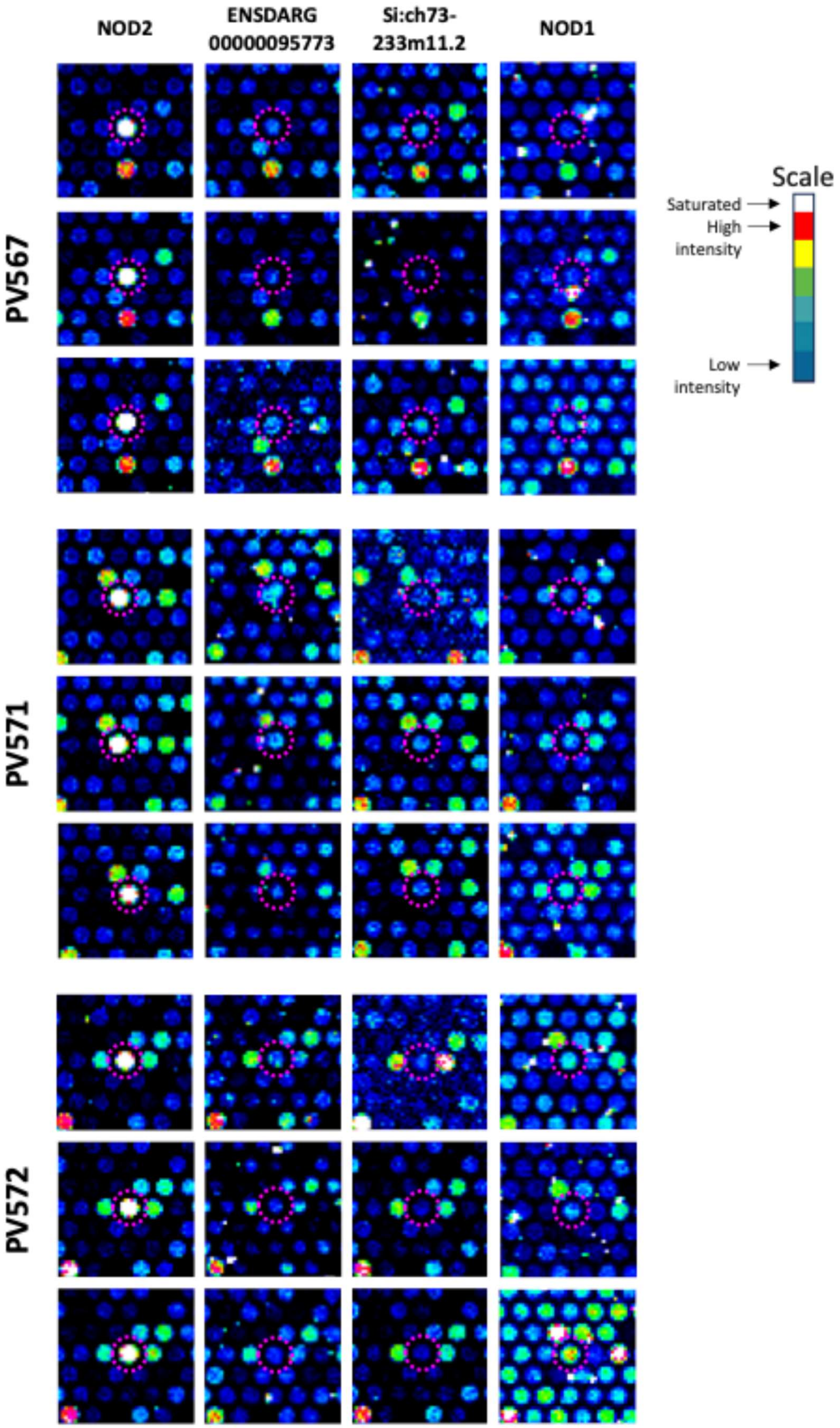
NOD2
ENSDARG 00000095773
Si:ch73-233m11.2
NOD1
PV567
PV571
PV572
Scale
Saturated
High intensity
Low intensity

**Figure S1. Representative images of APT-SNAP array binding.** Representative images of APT-SNAP array binding are shown for the three chosen aptamer candidates (PV567, PV571, and PV572) to the target protein NOD2, and three off-target proteins ENSDARG00000095773, si:ch73-233m11.2, and NOD1. The scale of fluorescence intensity (white are saturated pixels) observed is shown in the upper right legend. The aptamer feature of interest is in the center of each image and is indicated with a dashed magenta circle.

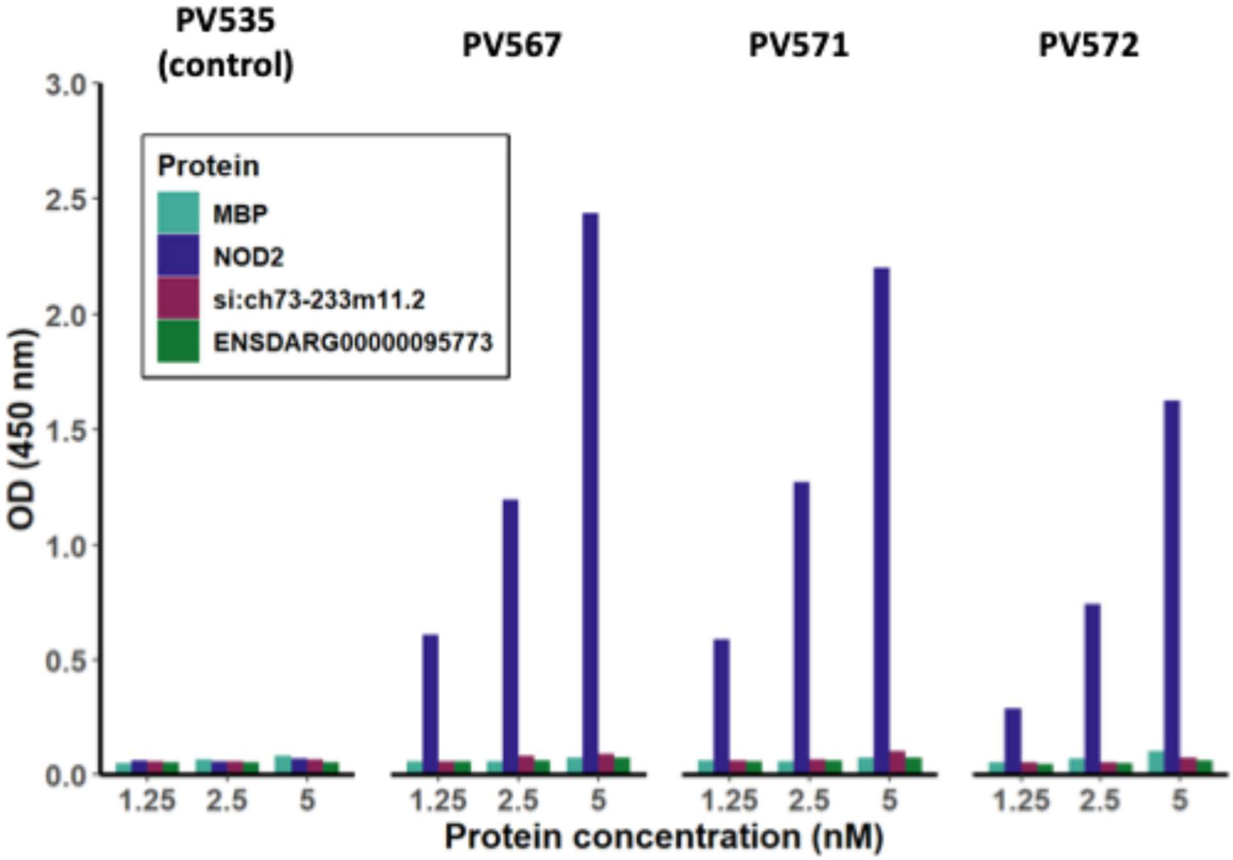


**Figure S2. NOD2 aptamer candidates demonstrate concentration dependence in ELASA format.** Binding of candidate aptamers to purified proteins was measured using enzyme-linked apta-sorbent assay (ELASA). Protein concentrations are indicated on the x-axis, and optical density at 450 nm on the y-axis.

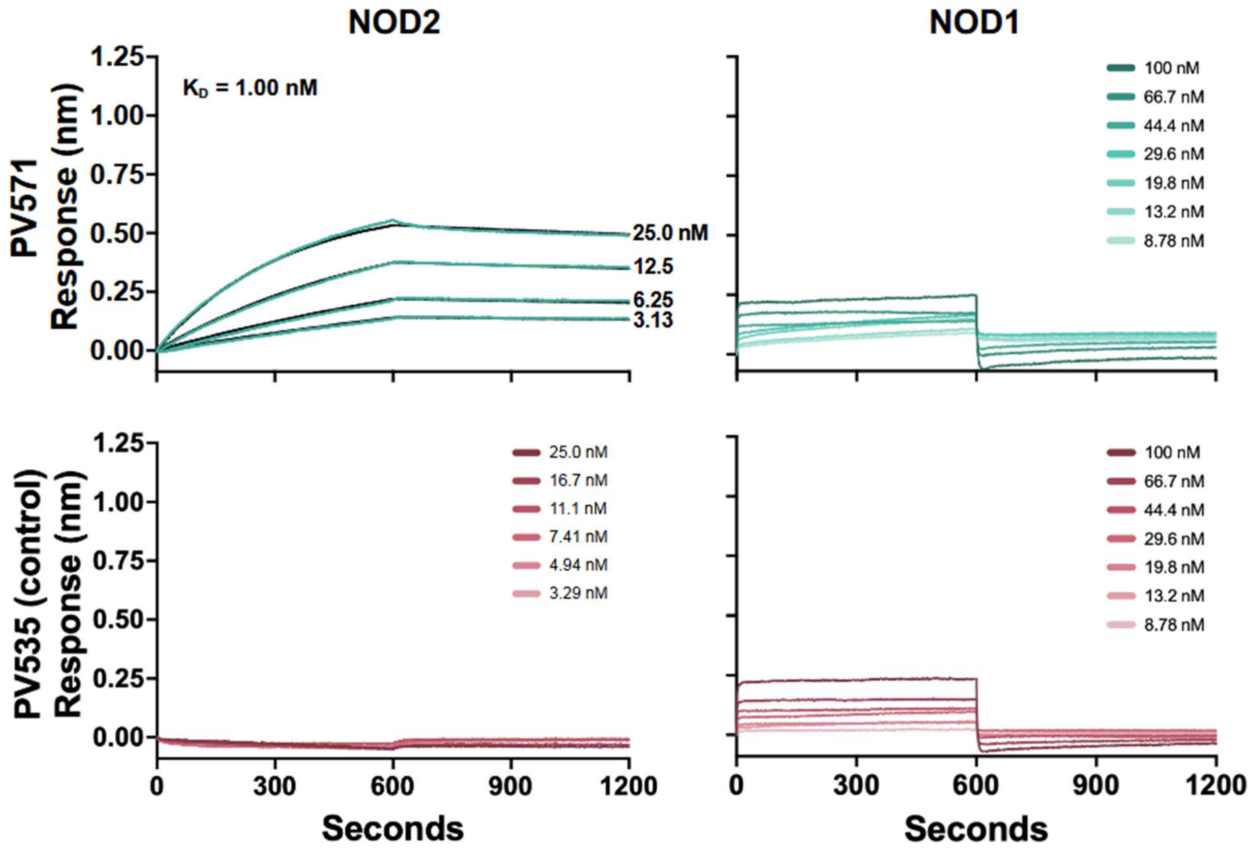


**Figure S3. Biolayer interferometry analysis of aptamer and control binding to NOD2 and NOD1.** Streptavidin biosensors were loaded with biotinylated DNA aptamers and exposed to purified protein at the range of indicated concentrations. Background-subtracted sensorgrams and global 1:1 kinetics fits (black) are shown. Aptamer PV571 binding (top) to NOD2 ($K_D$ = 1.00 nM) and no binding observed for off-target protein NOD1. Negative control PV535 (bottom) shows no binding to NOD2 or NOD1. Experiments were performed at 30 °C in Octet kinetics buffer supplemented with 5 mM $MgCl_2$.

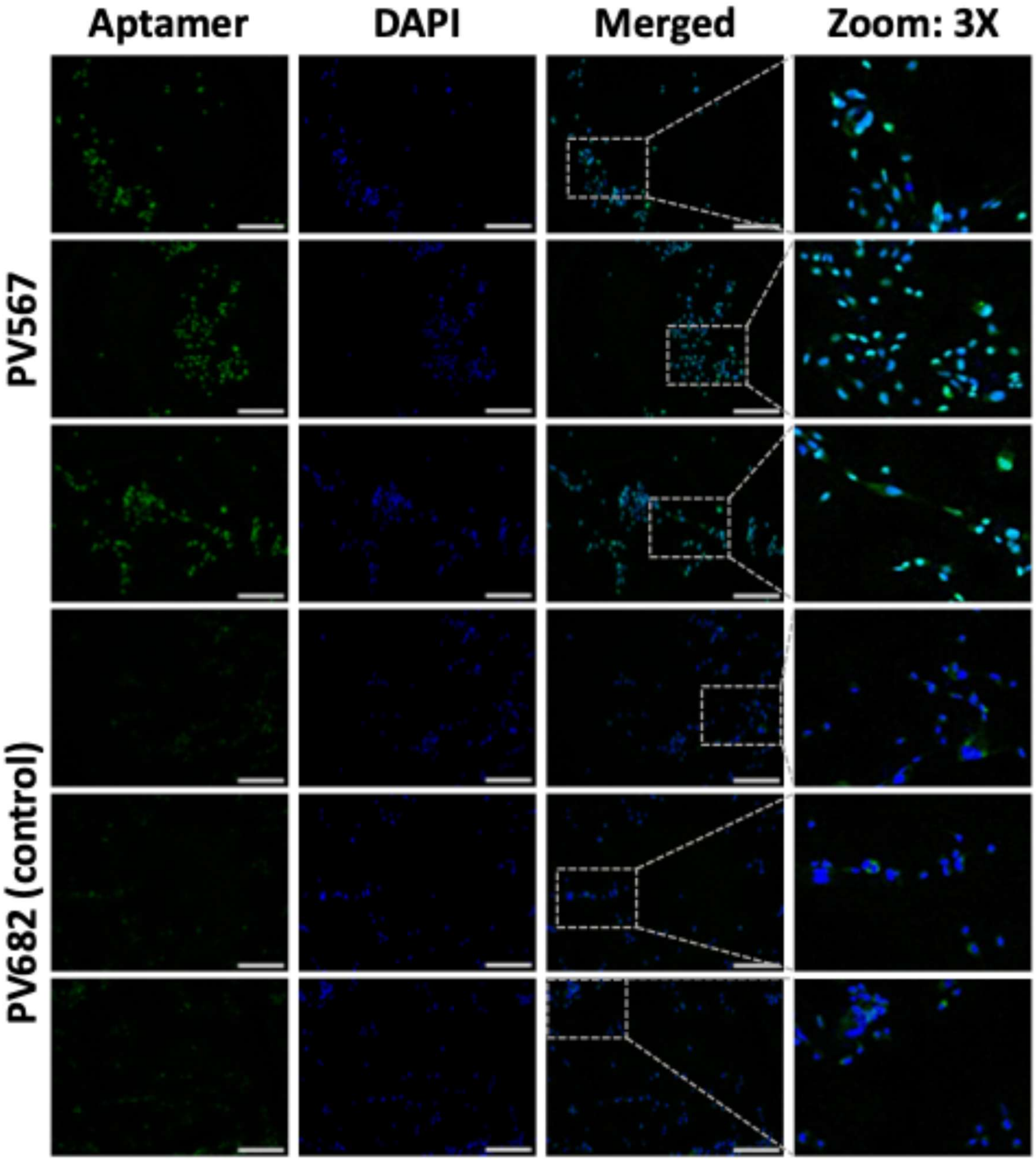


**Figure S4. Fluorescence microscopy detection of NOD2 candidate aptamers staining ZF4 cells.** Representative fluorescence microscopy images of cells stained with biotinylated aptamer (PV567) or negative control (PV682) followed by staining with streptavidin-Alexa Fluor™ 647 conjugate. Aptamer signal (green) and nuclear staining with DAPI (blue) are shown along with merged images and 3x zoomed regions. Scale bars are 200 µm. Images represent maximum contrast projections of 12 z-stack images acquired at 20 µm intervals. These and additional fields were used for quantification shown in Figure 7B.
